# Approximations to the expectations and variances of ratios of tree properties under the coalescent

**DOI:** 10.1101/2022.04.01.486796

**Authors:** Egor Lappo, Noah A Rosenberg

## Abstract

Properties of gene genealogies such as tree height (*H*), total branch length (*L*), total lengths of external (*E*) and internal (*I*) branches, mean length of basal branches (*B*), and the underlying coalescence times (*T*) can be used to study population-genetic processes and to develop statistical tests of population-genetic models. Uses of tree features in statistical tests often rely on predictions that depend on pairwise relationships among such features. For genealogies under the coalescent, we provide exact expressions for Taylor approximations to expected values and variances of ratios *X_n_*/*Y_n_*, for all 15 pairs among the variables {*H_n_*, *L_n_*, *E_n_*, *I_n_*, *B_n_*, *T_k_*}, considering *n* leaves and 2 ≤ *k* ≤ *n*. For expected values of the ratios, the approximations match closely with empirical simulation-based values. The approximations to the variances are not as accurate, but they generally match simulations in their trends as *n* increases. Although *E_n_* has expectation 2 and *H_n_* has expectation 2 in the limit as *n* → ∞, the approximation to the limiting expectation for *E_n_*/*H_n_* is not 1, instead equaling *π*^2^/3 – 2 ≈ 1.28987. The new approximations augment fundamental results in coalescent theory on the shapes of genealogical trees.

## 1 Introduction

Coalescent theory models random genealogies conditional on assumptions about the evolutionary process (Hein *et al*., 2005; Wakeley, 2009). In coalescent theory, a gene genealogy is a tree or network structure that represents a random draw from a coalescent model.

Genealogies in coalescent theory can be summarized using a variety of quantities. For example, for random tree-like genealogies with *n* lineages, the tree height *H_n_* records the sum of branch lengths on a path from leaves to the root, and the tree length *L_n_* sums all branch lengths in the tree. The total length *E_n_* of external branches sums over leaves the lengths of paths from leaves to their nearest internal nodes, and the total length of internal branches, *I_n_* = *L_n_* – *E_n_*, sums the lengths of all remaining branches.

Studies in coalescent theory have often investiated the properties of tree summaries conditional on assumptions of coalescent models, with the goal of understanding how shapes of the genealogies relate to processes such as population growth and migration (e.g. Slatkin & Hudson, 1991; Slatkin, 1996; Rosenberg & Feldman, 2002). Because mutations can be viewed as occurring conditionally on underlying genealogies (Hudson, 1990), features of genealogical shape affect the patterns of genetic variation produced by coalescent models that permit mutation. Thus, the understanding of summaries of tree shape predicted by coalescent models is a component of the interpretation of patterns of genetic variation in relation to evolutionary processes.

Initial results concerning summaries of genealogical shape focused on single quantities, producing results on expectations and variances of quantities such as *H_n_* and *L_n_* (Kingman, 1982; Hudson, 1983; Tajima, 1983). Subsequent studies examined the information that resides in the relationships between pairs of summaries; genetic variation statistics such as those of Tajima (1989) and Fu & Li (1993) can be viewed as assessing whether or not one aspect of a tree contains long branches in relation to another.

Recently, Arbisser *et al*. (2018) performed a detailed investigation of the relationship between *H_n_* and *L_n_* under coalescent models. They studied the mathematical relationship between these two quantities, computing under a standard coalescent model with a constant-sized population the covariance and correlation coefficient of *H_n_* and *L_n_*. Extending the work of Arbisser *et al*. (2018) on *H_n_* and *L_n_*, we (Alimpiev & Rosenberg, 2022) reported covariances and correlations for all pairs of variables among {*H_n_*, *L_n_*, *E_n_*, *I_n_*, *B_n_*, *T_k_*}, where *B_n_* is the mean of the lengths of the two basal branches of a genealogy and *T_k_* is the coalescence time from *k* to *k* – 1 lineages, 2 ≤ *k* ≤ *n*. Our compendium in Tables 1 and 2 of Alimpiev & Rosenberg (2022) summarizes pairwise relationships for several of the most commonly used features of coalecent tree shape, recording both new and previously known results.

**Table 1:** Definitions of random variables associated with various tree summaries. Here, *T_k_* is the coalescence time defined in Section 2.1, and 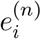 is the (random) length of the *i*th external branch of a tree with *n* leaves. We define *H_n_*, *L_n_*, and *E_n_* for *n* ≥ 2, *I_n_* for *n* ≥ 3, and *B_n_* for *n* ≥ 4.

| Variable | Definition |
| --- | --- |
| $H_n$ | $\sum_{k=2}^n T_k$ |
| $L_n$ | $\sum_{k=2}^n kT_k$ |
| $E_n$ | $\sum_{i=1}^n e_i^{(n)}$ |
| $I_n$ | $L_n - E_n$ |
| $B_n$ | $\frac{1}{2}T_2 + \left[ \sum_{j=3}^{n-1} \sum_{k=2}^j \frac{1}{j(j-1)} T_k \right] + \left( \sum_{k=2}^n \frac{1}{n-1} T_k \right)$ |

**Table 2:**
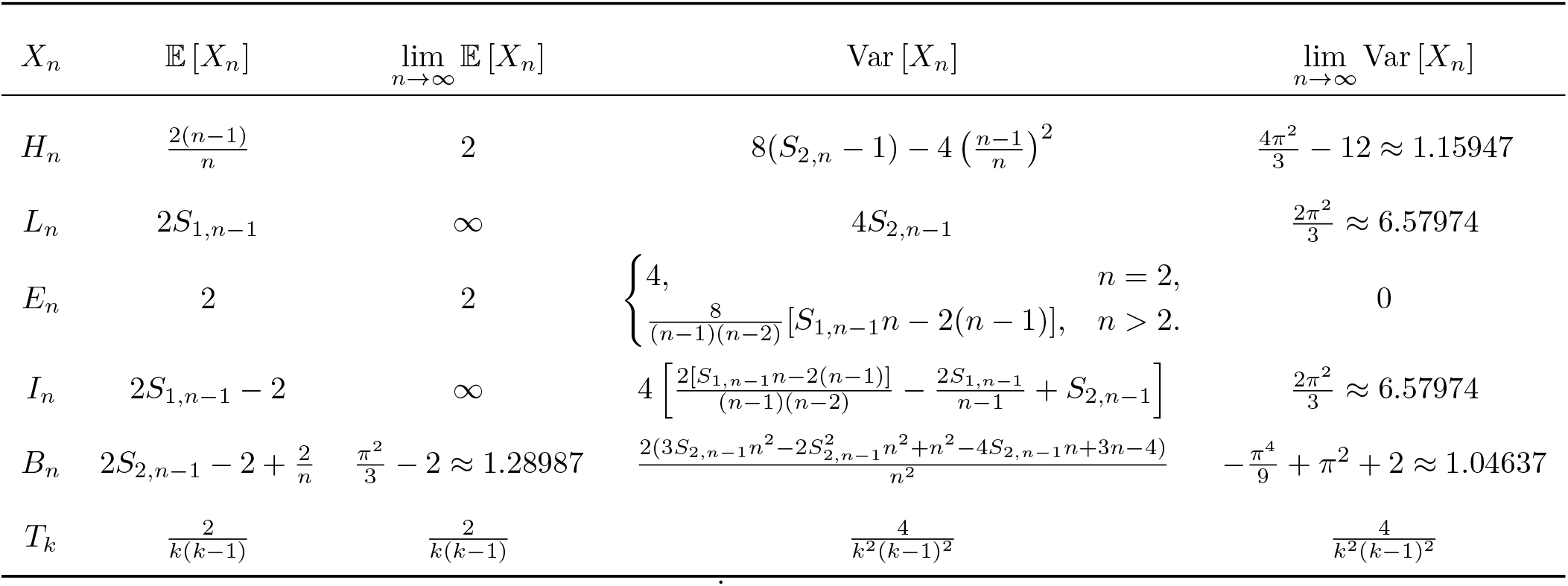
Expectations and variances of properties of tree branch lengths. These expressions can be found in Alimpiev & Rosenberg (2022)

| $X_n$ | $\mathbb{E}[X_n]$ | $\lim_{n \rightarrow \infty} \mathbb{E}[X_n]$ | $\text{Var}[X_n]$ | $\lim_{n \rightarrow \infty} \text{Var}[X_n]$ |
| --- | --- | --- | --- | --- |
| $H_n$ | $\frac{2(n-1)}{n}$ | 2 | $8(S_{2,n} - 1) - 4\left(\frac{n-1}{n}\right)^2$ | $\frac{4\pi^2}{3} - 12 \approx 1.15947$ |
| $L_n$ | $2S_{1,n-1}$ | ∞ | $4S_{2,n-1}$ | $\frac{2\pi^2}{3} \approx 6.57974$ |
| $E_n$ | 2 | 2 | $\begin{cases} 4, & n = 2, \\ \frac{8}{(n-1)(n-2)}[S_{1,n-1}n - 2(n-1)], & n > 2. \end{cases}$ | 0 |
| $I_n$ | $2S_{1,n-1} - 2$ | ∞ | $4\left[\frac{2[S_{1,n-1}n - 2(n-1)]}{(n-1)(n-2)} - \frac{2S_{1,n-1}}{n-1} + S_{2,n-1}\right]$ | $\frac{2\pi^2}{3} \approx 6.57974$ |
| $B_n$ | $2S_{2,n-1} - 2 + \frac{2}{n}$ | $\frac{\pi^2}{3} - 2 \approx 1.28987$ | $\frac{2(3S_{2,n-1}n^2 - 2S_{2,n-1}^2n^2 + n^2 - 4S_{2,n-1}n + 3n - 4)}{n^2}$ | $-\frac{\pi^4}{9} + \pi^2 + 2 \approx 1.04637$ |
| $T_k$ | $\frac{2}{k(k-1)}$ | $\frac{2}{k(k-1)}$ | $\frac{4}{k^2(k-1)^2}$ | $\frac{4}{k^2(k-1)^2}$ |

In addition to computing the covariance and correlation coefficient of *H_n_* and *L_n_*, Arbisser *et al*. (2018) also found approximations to the expectation and variance of the ratio *H_n_*/*L_n_* under the coalescent model. This ratio gives a summary of the joint distribution of *H_n_* and *L_n_* that characterizes the relative magnitudes of the variables—a feature not captured by their covariance or correlation. Arbisser *et al*. (2018) found that although the approximation to Var[*H_n_*/*L_n_*] differed noticeably from the exact value, as obtained by numerical integration and simulations of the coalescent model, the approximation to 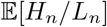 was quite accurate.

In this paper, we extend the work of Arbisser *et al*. (2018) to compute approximations to the expectations and variances for ratios of the 14 remaining pairs among {*H_n_*, *L_n_*, *E_n_*, *I_n_*, *B_n_*, *T_k_*}. The study performs for the expectation and variance of coalescent ratios an analogous extension of Arbisser *et al*. (2018) to that performed by Alimpiev & Rosenberg (2022) for the covariance and correlation coefficient.

## 2 Materials and methods

### 2.1 Tree variables

We work with a haploid population of constant size *N* that follows a standard coalescent model. Time is measured in units of *N* generations. In this section, we recall the definitions of the coalescence time *T_k_* and tree properties *H_n_*, *L_n_*, *E_n_*, *I_n_*, *B_n_* for sample size *n* ≥ 2 and 2 ≤ *k* ≤ *n*.

*T_k_* is defined to be a random variable representing the time to coalescence of *k* to *k* – 1 lineages, for 2 ≤ *k* ≤ *n*. Variable *T_k_* has exponential probability density function

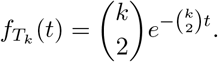

The expectation and variance of *T_k_* are

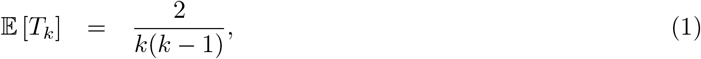

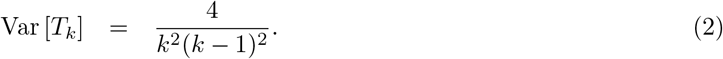

The tree properties *H_n_*, *L_n_*, *E_n_*, *I_n_*, and *B_n_* are defined in terms of the *T_k_*. Visual depictions of these properties appear in Figure 1, and mathematical definitions of these quantities appear in Table 1.

**Figure 1:**
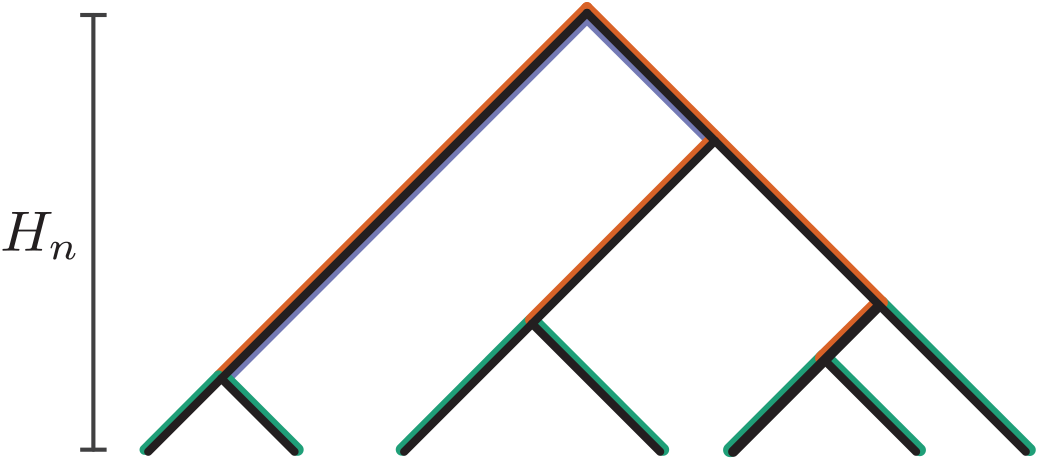
Properties of genealogical trees. The tree height is *H_n_*. The sum of the lengths of all branches is *L_n_*. External branches have total length *E_n_* (green). Internal branches have total length *I_n_* (red). Basal branches have mean length *B_n_* (purple).

We define 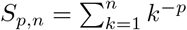 as a useful shorthand. The limit lim_*n*→∞_ *S_p,n_* = *S*_*p*,∞_ is the Riemann zeta function, usually denoted *ζ*(*p*). In particular, *S*_1,∞_ diverges, *S*_2,∞_ = *π*^2^/6 ≈ 1.64493, and *S*_3,∞_ is Apéry’s constant, approximately 1.20206.

### 2.2 Taylor approximations to expectations and variances of ratios

To compute approximate expressions for expected values and variances of the ratios of various tree properties, we rely on Taylor approximations. In particular, consider random variables *X* and *Y* with 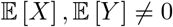. For the expectation, we have (second-order) approximation (Elandt-Johnson & Johnson, 1999, eq. 3.88)

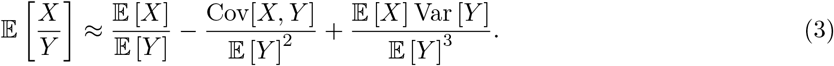

For the variance, we have (first-order) approximation (Stuart & Ord, 1994, eq. 10.17)

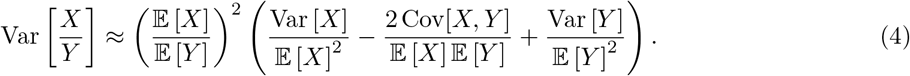

We use 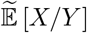 and 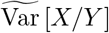 to denote approximations from eqs. 3 and 4. For both the expectation and the variance, we also take the *n* → ∞ limit of the approximations.

### 2.3 Exact expectations, variances, and covariances of tree properties

Expected values and variances of variables *H_n_*, *L_n_*, *E_n_*, *I_n_*, *B_n_*, and *T_k_* that are used in eqs. 3 and 4 are known, in many cases, from the earliest studies in coalescent theory (Fu & Li, 1993; Durrett, 2008; Wakeley, 2009). We summarize these expectations and variances in Table 2.

The covariances compiled by Alimpiev & Rosenberg (2022) appear in Table 3. In the case of pairs (*E_n_*, *B_n_*) and (*I_n_*, *B_n_*), the covariances are approximate, as described by Alimpiev & Rosenberg (2022).

**Table 3:** Covariances of pairs of variables that summarize genealogical trees. For pairs involving *E_n_* or *I_n_*, expressions apply for *n* ≥ 3; expressions involving *B_n_* apply for *n* ≥ 4. The expressions can be found in Alimpiev & Rosenberg (2022).

| $(X_n, Y_n)$ | $\text{Cov}[X_n, Y_n]$ | $\lim_{n \rightarrow \infty} \text{Cov}[X_n, Y_n]$ |
| --- | --- | --- |
| $H_n, T_k$ | $\frac{4}{k^2(k-1)^2}$ | $\frac{4}{k^2(k-1)^2}$ |
| $H_n, L_n$ | $4S_{2,n-1} - 4 + \frac{4}{n}$ | $\frac{2\pi^2}{3} - 4 \approx 2.57974$ |
| $H_n, E_n$ | $\frac{4}{n}$ | 0 |
| $H_n, I_n$ | $4S_{2,n-1} - 4$ | $\frac{2\pi^2}{3} - 4 \approx 2.57974$ |
| $H_n, B_n$ | $\frac{4[S_{3,n-1}n^2 - 3S_{2,n-1}n^2 + (4n+1)(n-1)]}{n^2}$ | $-2\pi^2 + 4\zeta(3) + 16 \approx 1.06902$ |
| $L_n, T_k$ | $\frac{4}{k(k-1)^2}$ | $\frac{4}{k(k-1)^2}$ |
| $L_n, E_n$ | $\frac{4S_{1,n-1}}{n-1}$ | 0 |
| $L_n, I_n$ | $4S_{2,n-1} - \frac{4S_{1,n-1}}{n-1}$ | $\frac{2\pi^2}{3} \approx 6.57974$ |
| $L_n, B_n$ | $\frac{4[S_{3,n-1}n - S_{2,n-1}n + n-1]}{n}$ | $-\frac{2\pi^2}{3} + 4\zeta(3) + 4 \approx 2.22849$ |
| $E_n, T_k$ | $\frac{4}{k(k-1)(n-1)}$ | 0 |
| $E_n, I_n$ | $\frac{4S_{1,n-1}}{n-1} - \frac{8S_{1,n-1}n}{(n-1)(n-2)} + \frac{16}{n-2}$ | 0 |
| $E_n, B_n$ | $\frac{4(S_{2,n-1}n - n + 1)}{n(n-1)}$ | 0 |
| $I_n, T_k$ | $\frac{4(n-k)}{k(k-1)^2(n-1)}$ | $\frac{4}{k(k-1)^2}$ |
| $I_n, B_n$ | $\frac{4(S_{3,n-1}n - S_{2,n-1}n + n - S_{3,n-1} - 1)}{n-1}$ | $-\frac{2\pi^2}{3} + 4\zeta(3) + 4 \approx 2.22849$ |
| $B_n, T_k$ | $\frac{4}{k^2(k-1)^3}$ | $\frac{4}{k^2(k-1)^3}$ |

### 2.4 Evaluating the approximations

For each of 15 pairs of random variables, considering *H_n_*, *L_n_*, *E_n_*, *I_n_*, and *B_n_* as well as *T_k_*, we substitute expressions from Tables 2 and 3 into eqs. 3 and 4 to obtain approximate expectations and variances for ratios of pairs of variables. For each pair, we choose one variable for the numerator and the other for the denominator; approximate expectations and variances for the reciprocals can be obtained similarly. We present the approximations in Tables 4 and 5, and we plot the approximations in Figures 2–5.

**Figure 2:**
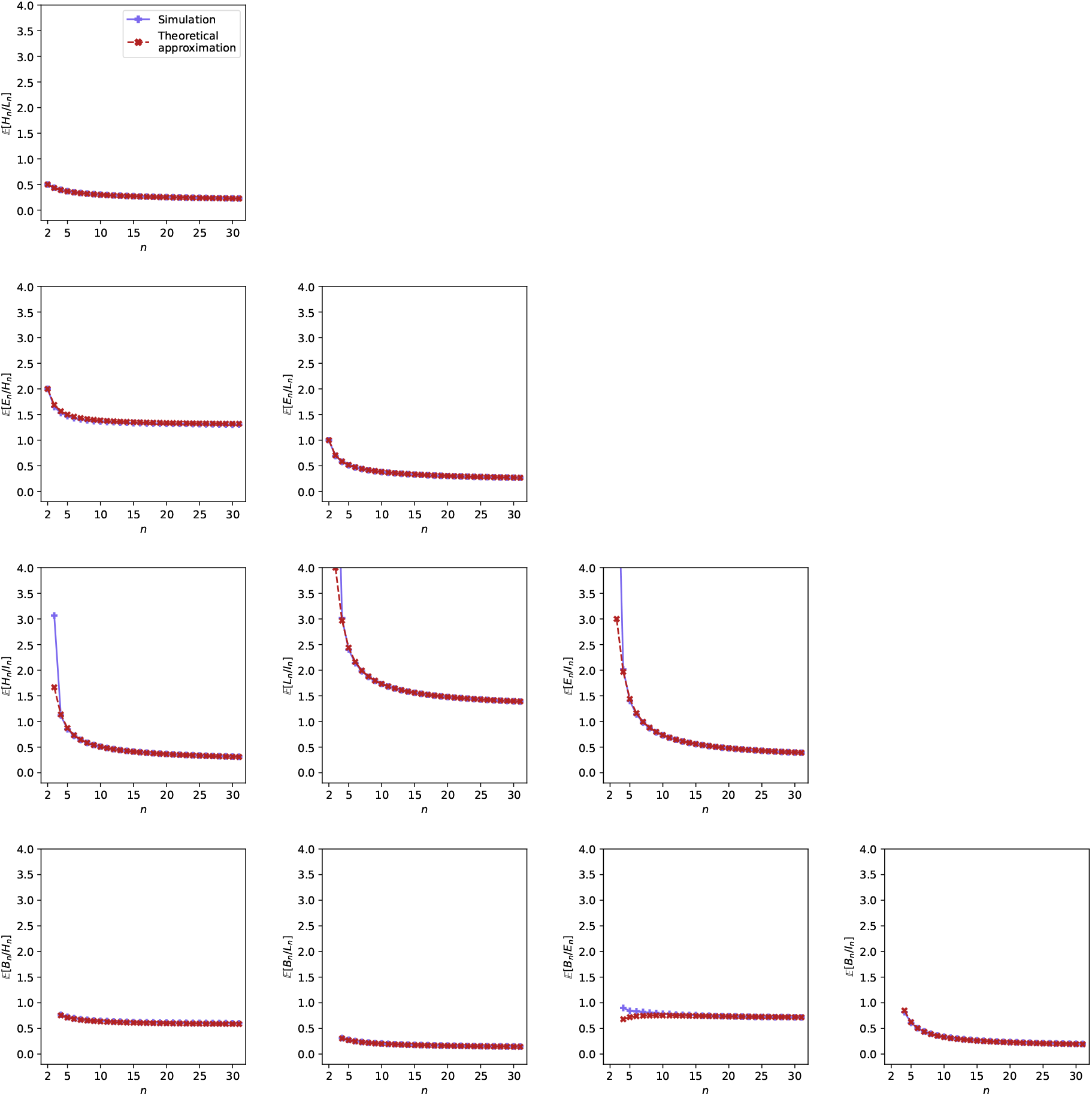
Simulated and theoretical approximations of expectations of ratios of pairs of variables, plotted as functions of sample size *n*. Expressions for theoretical values are taken from Table 4.

**Figure 3:**
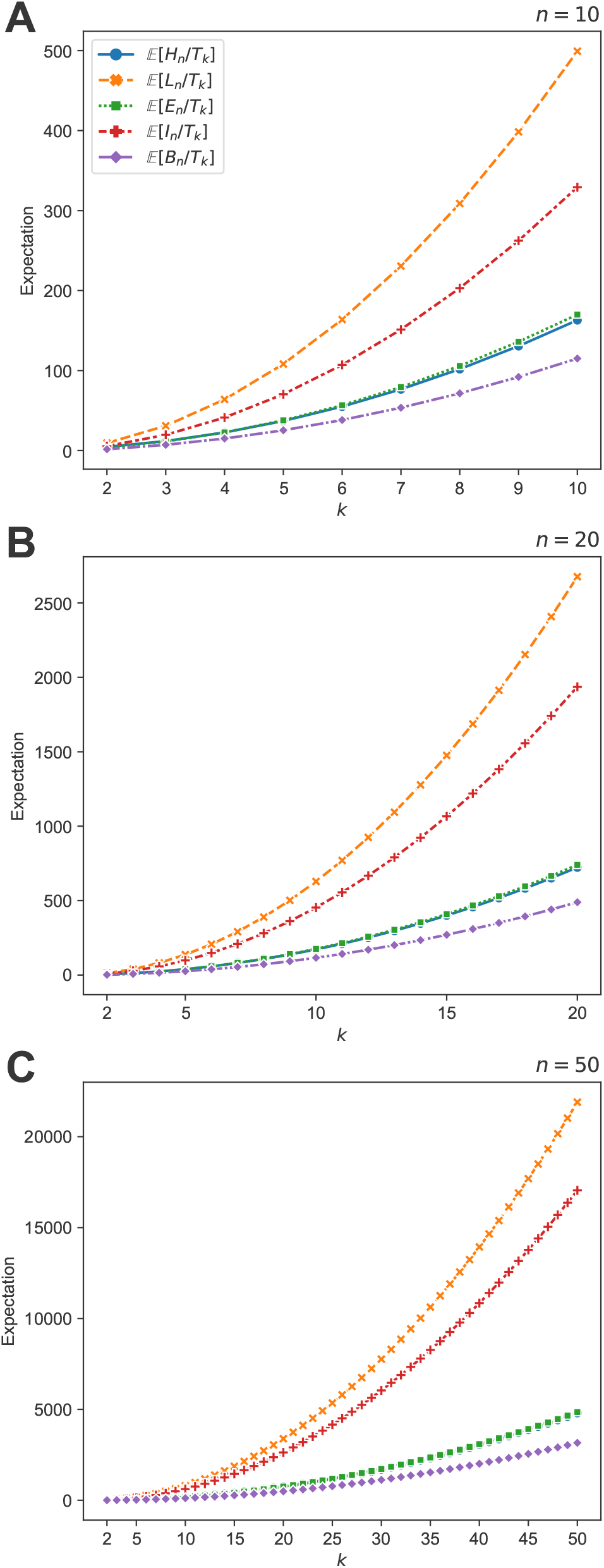
Theoretical approximations 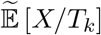 for variables *X* in {*H_n_*, *L_n_*, *E_n_*, *I_n_*, *B_n_*}, plotted as functions of *k* for *n* = 10, *n* = 20, and *n* = 50. The expressions plotted are taken from Table 4.

**Figure 4:**
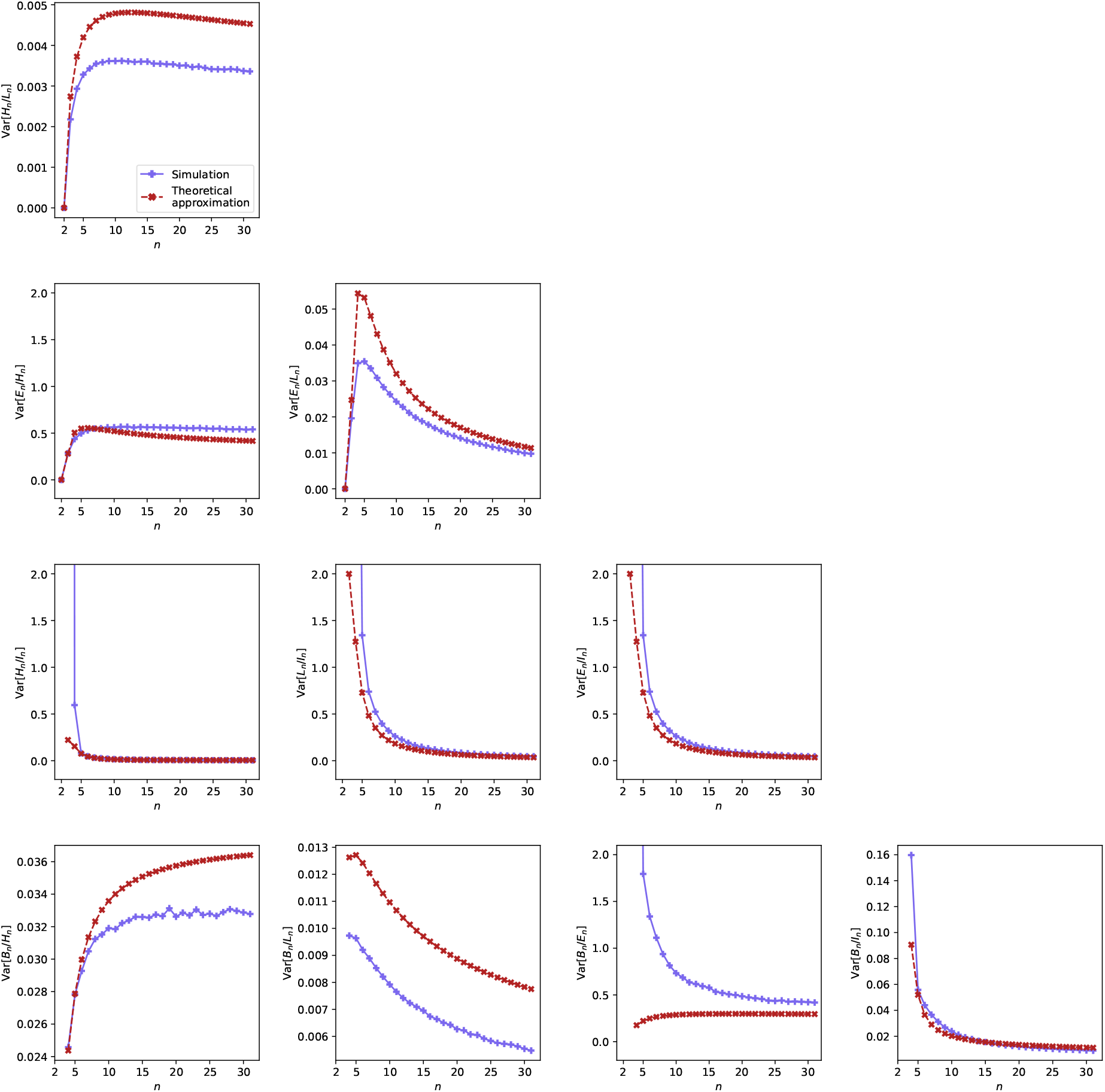
Simulated and theoretical approximations of variances of ratios of pairs of variables, plotted as functions of sample size *n*. Expressions for theoretical values are taken from Table 5.

**Figure 5:**
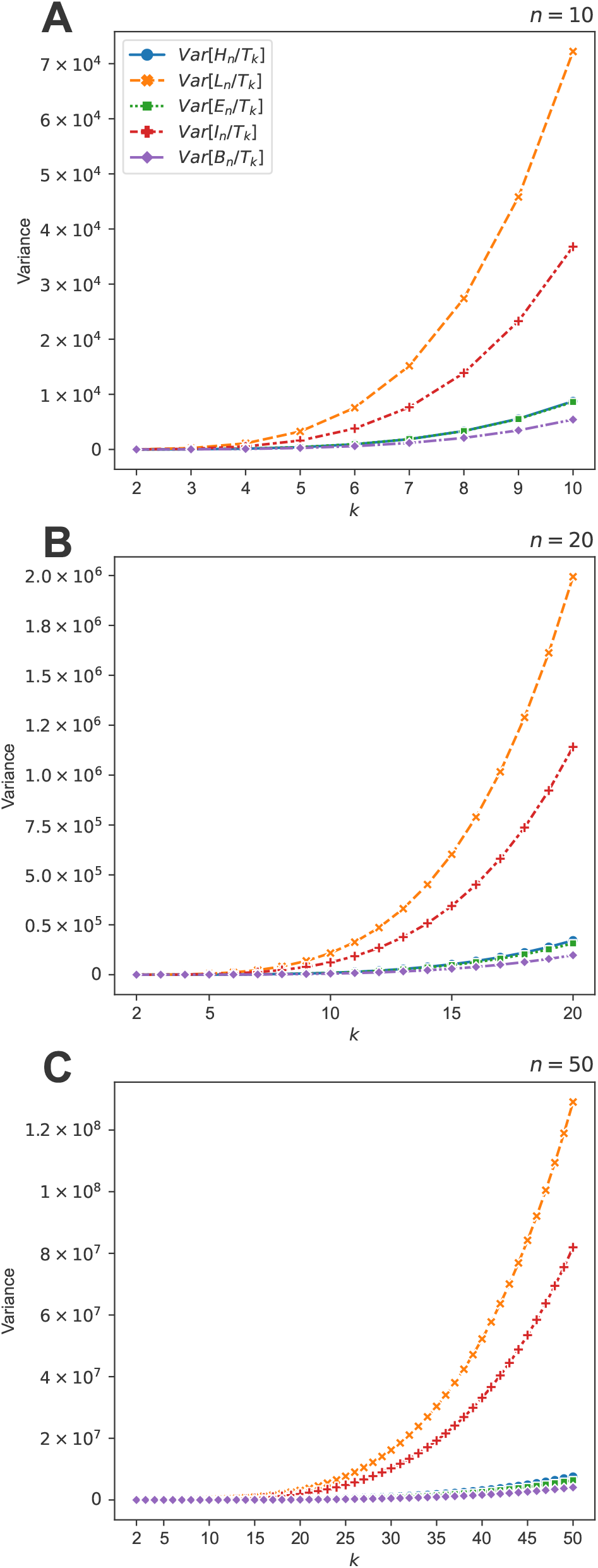
Theoretical approximations 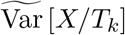 for variables *X* in {*H_n_*, *L_n_*, *E_n_*, *I_n_*, *B_n_*}, plotted as functions of *k* for *n* = 10, *n* = 20, and *n* = 50. The expressions plotted are taken from Table 5.

**Table 4:**
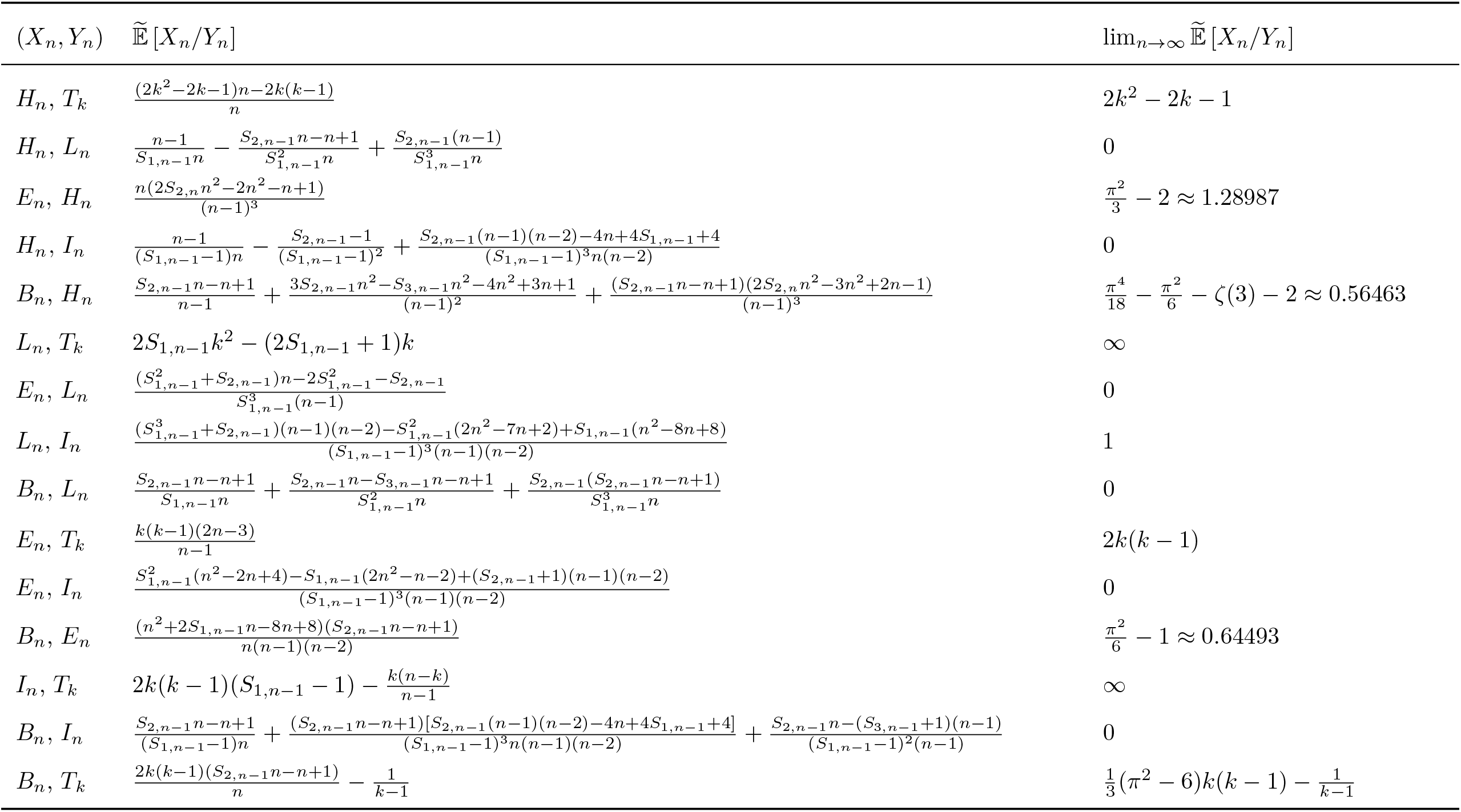
Approximations to expectations of ratios of pairs of variables. Expressions involving *E_n_* or *I_n_* apply for *n* ≥ 3; expressions involving *B_n_* apply for *n* ≥ 4. The value for (*H_n_*, *L_n_*) follows eq. 15 of Arbisser *et al*. (2018). The expressions are obtained using eq. 3 and Tables 2 and 3.

**Table 5:**
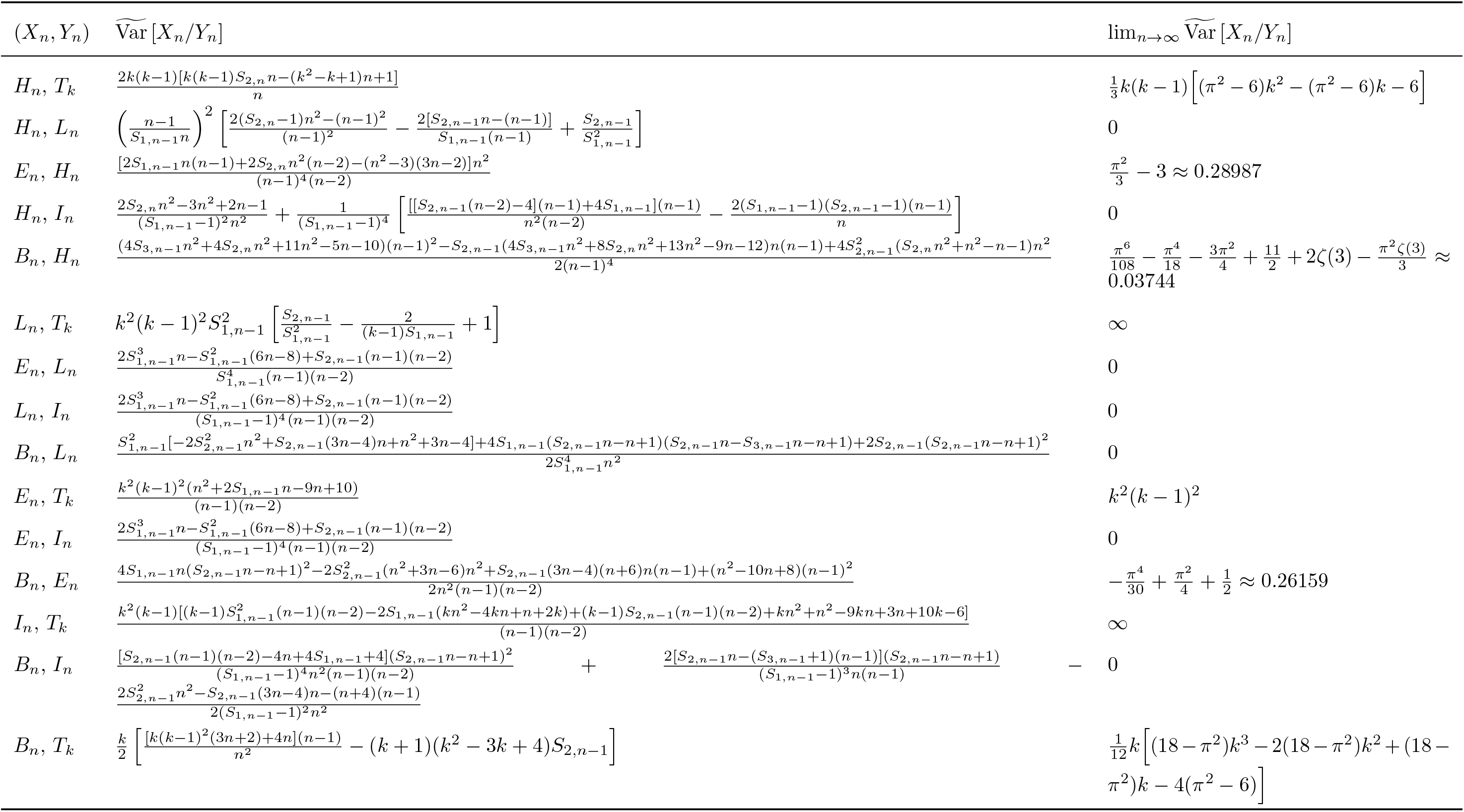
Approximations to variances of ratios of pairs of variables. Expressions involving *E_n_* or *I_n_* apply for *n* ≥ 3; expressions involving *B_n_* apply for *n* ≥ 4. The value for (*H_n_*, *L_n_*) follows eq. 18 of Arbisser *et al*. (2018). The expressions are obtained using eq. 4 and Tables 2 and 3.

For pairs (*X_n_*, *Y_n_*), we simulate the values of 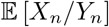 and Var [*X_n_*/*Y_n_*] under the coalescent model using using `ms` (Hudson, 2002), performing 100,000 replicate simulations for each tree size *n* = 2, 3,…, 50. We plot the simulated values alongside the approximate values from Tables 4 and 5 in Figures 2 and 4.

## 3 Results

### 3.1 Expectations of the ratios

The approximate expected values in Table 4, as approximations of ratios, have the form of rational functions. As *n* grows, the approximate expectations of *H_n_*/*L_n_*, *H_n_*/*I_n_*, *E_n_*/*L_n_*, *E_n_*/*I_n_*, *B_n_*/*L_n_*, and *B_n_*/*I_n_* approach 0. This behavior is sensible when considering the properties of the coalescent model: in the numerators, *E_n_* has expectation 2 and 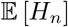 and 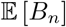 have bounded expectation in the limit as *n* → ∞; in the denominators, *L_n_* and *I_n_* have expectations that grow without bound (Table 2). Similarly, approximate expectations of ratios *L_n_*/*T_k_* or *I_n_*/*T_k_* with *L_n_* and *I_n_* in the numerator and *T_k_* in the denominator grow to infinity as *n* increases. The approximation to 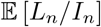 approaches 1 in the limit as *n* → ∞: as the number of leaves in the tree grows, internal branches occupy an increasingly large fraction of the total branch length.

For pairs of variables that both have finite expectation, the approximate expectations of their associated ratios—*H_n_*/*T_k_*, *E_n_*/*H_n_*, *E_n_*/*T_k_*, *B_n_*/*H_n_*, *B_n_*/*E_n_*, and *B_n_*/*T_k_*—also approach finite values in the limit as *n* → ∞. It is interesting to observe that although 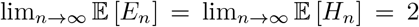 (Table 2), 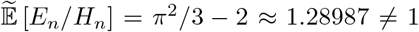. In other words, although expectations of the individual variables approach the same value, we expect *E_n_*/*H_n_* to be somewhat larger than 1 on average.

For each of the 10 pairs of variables among {*H_n_*, *L_n_*, *E_n_*, *I_n_*, *B_n_*}, the approximate expectations from Table 4 are plotted in Figure 2 together with the simulated values. Although some divergences are present for small *n*, the approximate and simulated values match closely.

The approximate ratios involving T_k_ are shown in Figure 3 as functions of *k* for each of three values of *n*. *L_n_* is the fastest-growing variable according to the expression for its expecteation (Table 2), and the graph for 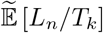 is topmost in all three plots. As expectations of *H_n_* and *E_n_* are close (Table 2), the graphs for 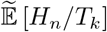 and 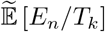 are close in Figure 3.

### 3.2 Variances of the ratios

The limits of approximations of variances of ratios are presented in Table 5. They behave similarly to the expectations in Table 4. Because *L_n_* and *I_n_* have expectations that grow without bound, for ratios *H_n_*/*L_n_*, *H_n_*/*I_n_*, *E_n_*/*L_n_*, *B_n_*/*L_n_*, *E_n_*/*I_n_*, *B_n_*/*I_n_*—with *L_n_* or *I_n_* in the denominator—the limits of the variance approximations are 0. As *n* grows, the denominators grow much faster than the numerators, and the values are therefore increasingly concentrated around 0. Hence, the variances also approach 0.

Because *L_n_* and *I_n_* are much larger than the coalescence times *T_k_*, approximations to variances of *L_n_*/*T_k_* and *I_n_*/*T_k_* diverge to infinity as *n* increases. Interestingly, however, the approximate variance of *L_n_*/*I_n_*, a ratio of two quantities with diverging expectations, approaches 0.

The variance approximations with finite nonzero limits are those for *H_n_*/*T_k_*, *E_n_*/*H_n_*, *E_n_*/*T_k_*, *B_n_*/*H_n_*, *B_n_*/*E_n_*, and *B_n_*/*T_k_*. All give ratios of two variables with finite expectation and variance as *n* → ∞ (Table 2).

Figure 4 shows the expressions from Table 5 together with the simulated values. Compared to the plots of expectations of ratios (Figure 2), differences between the simulated and approximate variances are prominent at small n. For the variances of *H_n_*/*L_n_*, *B_n_*/*H_n_*, and *B_n_*/*L_n_*, the simulated and approximate values differ substantially even as *n* increases. Because the theoretical value of Cov[*E_n_*, *B_n_*] that contributes to the approximate variance of *B_n_*/*E_n_* is itself an approximation, one of the larger differences between simulation and approximation occurs for the plot for 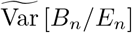.

Figure 5 shows variances of ratios involving *T_k_* for varying *k*, for each of three values of *n*. Qualitatively, the values for approximate variances behave similarly to expectations in Figure 3: in particular, the vertical placement of the curves follows the same order. Our approximations to the variance of *L_n_*/*T_k_* and *I_n_*/*T_k_* grow fastest, as the numerators are typically large and the expected value of the denominator *T_k_* decreases as *k* grows. Approximations to variances of *H_n_*/*T_k_*, *E_n_*/*T_k_*, and *B_n_*/*T_k_* all display much slower growth; for these quantities, the expectations of numerators of the ratios are bounded above by 2 for all *n*.

## 4 Discussion

In this paper, we have computed approximations to expected values and variances of ratios of various branch lengths under the standard coalescent model. We have considered all 15 possible pairs of variables among {*H_n_*, *L_n_*, *E_n_*, *I_n_*, *B_n_*, *T_k_*}. We have also assessed the accuracy of approximations to the expectation and variance by comparing them with values computed by simulation. We have observed that the approximate expressions behave in a way that matches mathematical intuition about the behavior of random variables associated with the branch lengths.

As *n* grows large, the ratios involving *L_n_* and *I_n_* have nearly identical behavior, explained by the fact that internal branches take up increasingly large fractions of the total branch length (Figures 2–5). In the limit as *n* → ∞, expectations of both *H_n_* and *E_n_* approach a constant value of 2 (Table 2), and Cov[*H_n_*, *E_n_*] approaches 0 (Table 3). However, we observed that 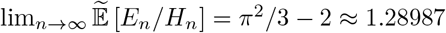 is not equal to 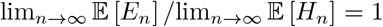. For the ratio *B_n_*/*E_n_*, the approximation aligns with the naive prediction, 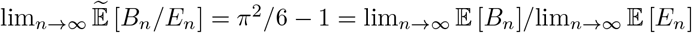, even though Cov[*E_n_*, *B_n_*] is also zero in the limit (Table 2). For *B_n_* and *H_n_*, which possess a high correlation, 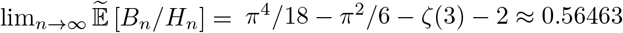, whereas 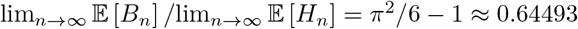.

Previously, we evaluated covariances correlation coefficients under the coalescent model for the pairs of variables that we consider here, obtaining exact covariances and correlations for 13 of 15 pairs and approximations for the other two. We obtained limiting expressions for these covariances and correlations as *n* → ∞. The approximate values that we have provided here for expectations and variances of ratios make use of these previous results concerning covariances, adding to the understanding of the properties of joint distributions of pairs of genealogical variables in coalescent theory.

Ratios between coalescent branch lengths have potential uses in understanding the effects of population processes on the shapes of evolutionary trees. In particular, tests that evaluate site-frequency spectra for agreement with predictions of coalescent models can often be viewed as assessing if specified random coalescent quantities equal a null value, such as 0 (Fu, 1995; Zeng *et al*., 2006; Achaz, 2009; Ferretti *et al*., 2010; Ronen *et al*., 2013; Ferretti *et al*., 2017). Such tests are often based on a recognition that two quantities have equal expectation and an expected difference of 0, rather than based on an analysis of a ratio of the two quantities, although several modeling studies do emphasize ratios (Slatkin, 1996; Uyenoyama, 1997; Rosenberg & Hirsh, 2003; Arbisser *et al*., 2018). The approximate expectation and variance can augment understanding of scenarios in which coalescent ratios are considered.

We have found that approximations for fixed *n* and in the limit as *n* → ∞ are quite accurate in predicting the expected values seen in coalescent simulations of the ratios (Figure 2). For the variances, the approximations are generally less accurate, although in most cases, the shapes of graphs of the approximations and simulated values are similar (Figure 4). Higher-order approximations than eq. 4 would potentially be required for improving the agreement between approximate and simulated values.

## Data availability

The `ms` command for simulations is `ms n 100000 -T`, where `n` is taken from {2, 3,…, 50} and gives the number of leaves of simulated trees.

## Acknowledgments

We acknowledge support from NIH grants R01 GM131404 and R01 HG005855 and NSF grant BCS-2116322.

